# Utilization of the repeated squat-stand model for studying the directional sensitivity of the cerebral pressure-flow relationship

**DOI:** 10.1101/2020.12.15.422722

**Authors:** Lawrence Labrecque, Jonathan D Smirl, Patrice Brassard

**Author notes:** **Address for correspondence:** Patrice Brassard, Ph.D., Department of Kinesiology, Faculty of Medicine, PEPS - Université Laval, 2300 rue de la Terrasse, room 0290-H, Québec (Qc) GIV OA6, Canada, **Email:**.

## Abstract

Hysteresis in the cerebral pressure-flow relationship describes the superior ability of the cerebrovasculature to buffer cerebral blood flow changes when mean arterial pressure (MAP) acutely increases compared to when MAP acutely decreases. This phenomenon can be evaluated by comparing the change in middle cerebral artery mean blood velocity (MCAv) per change in MAP during either acute increases or decreases in MAP induced by repeated squat-stands (RSS). However, no real baseline can be employed for this particular protocol as there is no true stable reference point. Herein, we characterized a novel metric using the greatest MAP oscillations induced by RSS without using an independent baseline value and adjusted for time intervals (ΔMCAv_T_/ΔMAP_T_). We also examined whether this metric during each RSS transition were comparable between each other over a 5-min period. ΔMCAv_T_/ΔMAP_T_ was calculated using the minimum to maximum MCAv and MAP for each RSS performed at 0.05 Hz and 0.10 Hz. We compared averaged ΔMCAv_T_/ΔMAP_T_ during MAP increases and decreases in 74 healthy subjects [9 women; 32 ± 13 years]. ΔMCAv_T_/ΔMAP_T_ was lower for MAP increases than MAP decreases at 0.10 Hz RSS only (0.91 ± 0.34 vs. 1.01 ± 0.44 cm·s^-1^/mmHg; p = 0.0013). For both frequency and MAP direction, time during RSS had no effect on ΔMCAv_T_/ΔMAP_T_. This novel analytical method supports the use of the RSS model to evaluate the directional sensitivity of the pressure-flow relationship. These results contribute to the importance of considering the direction of MAP changes when evaluating dynamic cerebral autoregulation.

**News & Noteworthy:** Repeated squat-stand maneuvers are able to examine the directional sensitivity of the cerebral pressure-flow relationship. These maneuvers induce stable physiological cyclic changes where brain blood flow changes with blood pressure increases are buffered more than decreases. These results highlight the importance of considering directional blood pressure changes within cerebral autoregulation.

## Introduction

Dynamic cerebral autoregulation (dCA) defines the ability of the cerebrovasculature to react to rapid blood pressure changes. Evidence suggests asymmetrical responses of cerebral blood flow (CBF) to transient changes in mean arterial pressure (MAP). Specifically, the augmentation in CBF is attenuated when MAP acutely increases compared to the decline in CBF when MAP acutely decreases (1, 8, 27, 36). Since the brain is located in the skull, an inflexible structure, this phenomenon is thought to serve as a protective mechanism against blood pressure surges (39). This notion has been previously highlighted as the cerebrovasculature is more efficient at buffering changes in blood pressure associated with the systolic aspect of the cardiac cycle (11, 32).

Initially, the above-mentioned asymmetrical responses of CBF to acute changes in MAP have been reported using cyclic inflation and deflation of thigh cuffs in patients with head injury (1) and pharmacological interventions in healthy participants (36). It is important to ensure the hysteresis-like pattern of the cerebral pressure-flow relationship exists during non-pharmacologically and physiologically induced acute increases and decreases in MAP in health and disease. Accordingly, the development of a valid non-pharmacological model to examine the directional sensitivity of the cerebral pressure-flow relationship is imperative.

Unfortunately, the previously used thigh cuff deflation technique may not represent the best model to investigate the directional sensitivity of the cerebral pressure-flow relationship, considering the poor reproducibility associated with dCA metrics quantified with this technique (reviewed in (29)). In the cerebrovascular research community, a repeated squat-stand protocol of 5 min is recognized as a reproducible method for inducing large blood pressure oscillations aimed at improving the linear interpretability of the transfer function metrics (29). This 5-min period of repeated squat-stands has been commonly used to examine the relationship between cerebral blood velocity (CBV) and MAP in the frequency domain at two different repeated squat-stands frequencies (0.05 and 0.10 Hz) with the transfer function analysis, an analytical method which does not consider the direction of MAP (15). These two squatting frequencies are utilized since they are included in the frequency bands where dCA is thought to have the most important influence on the cerebral pressure-flow dynamics (historically, these frequency bands being 0.02-0.07 Hz for the very low frequency and 0.07-0.20 for the low frequency) (15). Also, these two frequencies are most prevalent in the literature (7, 10-12, 14, 18, 21, 22, 25, 28-31). Interestingly, the repeated squat-stand maneuver induces a cyclic physiological stress that can also be utilized to evaluate the CBV-MAP relationship in the time domain (6, 8, 27). When comparing the relative change in middle cerebral artery mean blood velocity (MCAv) per relative change in MAP during repeated squat-stands, our team previously demonstrated the directional sensitivity of the cerebral pressure-flow relationship (8). Using the autoregulation index (ARI), Panerai et al. demonstrated a better autoregulatory capacity during the squatting phase (transient increase in MAP) than the standing phase (transient decrease in MAP) (27). These results support the usefulness of analyzing MAP oscillations induced by the repeated squat-stand model in the time domain to evaluate the directional sensitivity of the cerebral pressure-flow relationship. Furthermore, the 5-min repeated squat-stand model allows for the average of several induced responses. Thus, providing a more reliable estimate of the physiological response than the utilization of only one transient response for each MAP direction as had previous be performed with the rate-of-regulation (2) and the autoregulatory index (34) measures. Additionally, we need to assume the metric used to characterize the hysteresis-like pattern during each squat-stand repetition is comparable over the 5 min, which is presently unknown.

In our previous analysis, we characterized the directional sensitivity of the cerebral pressure-flow relationship using the change in MCAv in response to transient increases versus decreases in MAP induced by repeated squat-stands. Specifically, we averaged the percent change in MCAv (from baseline) per percent change in MAP (from baseline) (%ΔMCAv/%ΔMAP) calculated for each squat-stand executed over a period of 5 min at the frequency of interest (0.05 and 0.10 Hz). These relative changes were calculated from a seated baseline obtained prior-to the repeated squat-stand maneuver (8). However, the utilization of a separate resting period as the baseline value has been recently criticized (6). Barnes et al. argued the magnitude of MAP oscillations induced by repeated squat-stands is greater when utilizing resting seated MAP as a reference value compared to a resting standing MAP (6). In fact, they reasoned that the decrease in MAP from a resting seated position to the minimum reached during a stand, is greater and would explain the hysteresis pattern. These authors suggested using a reference value in the standing position immediately before the repeated squat-stand protocol would be more valid.

While we agree calculating a baseline value from a separate control recording may be misleading, the utilization of a reference value from a prolonged standing position immediately before the beginning of the repeated squat-stand maneuver can also be problematic. In this case, the increase in MAP from a resting quiet-stance position to the maximum reached during a squat would be greater than the MAP decrease and balance the difference between both. In addition, a prolonged period of standing, necessary to reach a steady state of measured variables, has the potential to lead to orthostatic hypotension in some individuals. In light of these issues, we consider there is no real baseline we can employ for this particular protocol as there is not a truly stable physiological reference point we can compare the changes during the squat-stand maneuvers to.

A notion we did not consider in our previous study was the time intervals when the transition of MCAv and MAP took place, which represents an additional issue with this work (8). There are delays in physiological responses and regulating mechanisms vary whether changes in MCAv and MAP are increasing or decreasing (1). Therefore, we cannot assume repeated squat-stands induce MCAv and MAP changes of comparable time intervals when MAP acutely decreases and increases via repeated squat-stands induced at 0.05 Hz or 0.10 Hz. Thus, upward (from minimum to maximum) and downward (from maximum to minimum) changes in MCAv/MAP of similar amplitudes, but occurring across different time intervals, would indicate vessels are reacting differently according to the direction of the MAP changes. We also believe taking into consideration time intervals when MCAv and MAP changes are taking place would be more appropriate in order to compare our metric between RSS frequencies (0.05 vs. 0.10 Hz).

The prevalent modeling techniques utilized for dCA quantification with spontaneous or driven blood pressure oscillations do not take the directionality of blood pressure changes into consideration (i.e. autoregulation index, transfer function analysis of spontaneous and forced blood pressure oscillations). In other words, these approaches assume the cerebrovasculature reacts similarly when MAP acutely increases or decreases. Confirmation of asymmetrical sensitivity of the cerebral pressure-flow relationship using the repeated squat-stand model thus has the potential to refine and optimize the current methods of dCA assessment (15). If it becomes clear this hysteresis-like pattern of the cerebral pressure-flow relationship is a common characteristic of the healthy cerebrovasculature, which could be influenced by physiological and pathological conditions, there will be a need for a global change in the assessment of dCA in order to take MAP direction into account. However, the optimal physiological approach to study this phenomenon remains to be elucidated.

For these reasons, the primary aim of this study was to refine a metric using the largest concurrent MAP oscillations induced by the repeated squat-stand maneuver without using an independent baseline value (either seated or standing). Accordingly, we calculated this metric by using the minimum to maximum MCAv and MAP (in response to squatting) compared to the maximum to minimum MCAv and MAP (in response to standing) for each squat-stand transition taking time intervals when the MCAv and MAP transition took place into account. The secondary aim of this study was to examine whether the novel metric during each squat-stand transition at both 0.05 and 0.10 Hz were comparable between each other over the 5-min period. We hypothesized the metric would be lower when MAP acutely increases compared to when MAP acutely decreases, and this 5-min repeated squat-stand model is an adequate protocol for the characterization of the metric since it will be comparable between all repeated squat-stand transitions.

## Material and methods

### Participants

Seventy-four participants were included in this analysis. Participants had previously been involved in research studies at one of two institutions (Université Laval, Canada; University of British Columbia, Canada). All participants provided informed written consent before participating to these studies. The original research studies were approved by the *Comité d’éthique de la recherche de l’Institut universitaire de cardiologie et de pneumologie de Québec* (CER : 20869 and 21180) and the University of British Columbia Clinical Review Ethical Board (CER: H11-02576, H11-03287) according to the principles established by the Declaration of Helsinki (except for registration in a database). Since the current study included a secondary analysis from previous institutional review board approved studies and the use of de-identified data sharing, this study did not require additional institutional review board review. All the participants were healthy, free from medical condition and had no history of cardiorespiratory or cerebrovascular disease. Apart from oral contraceptives in some women, participants were not taking any blood pressure-altering medication. Women included in the study were either taking oral contraceptive continuously for more than a year (n=2), wearing an intrauterine device (n=1) or were tested during menses or the early follicular phase (day 1 to 10) of their menstrual cycle (n=6).

### Experimental protocol

The experimental protocol has been described previously (8). Participants arrived fasted at the laboratory around 7AM and had a standardized snack (granola bar and juice). An initial 5-min seated baseline recording was made for all participants for baseline characteristics. Following a 5-min standing rest, the participants performed repeated squat-stands for a period of at least 5 min at a frequency of 0.05 Hz (10-s squat, 10-s standing) and 0.10 Hz (5-s squat, 5-s standing). As previously mentioned, these frequencies were selected since they are within the range where dCA is thought to have its greatest influence on the cerebral pressure-flow dynamics (40). Familiarization with the squat-stand movement was done by mirroring the experimenter before the experimentation. During recording, the experimenter ensured the transition speed and depth was similar throughout the squat-stand repetition protocol for each participant.

### Instrumentation

Maximal oxygen consumption (VO_2max_) was determined during a preliminary visit in 53 participants. All these participants performed a graded ramp exercise test up to exhaustion on an electromagnetically braked upright cycling ergometer. After 1 min of unloaded pedaling, the workload increased progressively (20-30 W/min) until volitional exhaustion. A gas exchange analysis system, combining fast responding oxygen and carbon dioxide sensors and a pneumotach (Breezesuite, MedGraphics Corp., MN, USA; Ultima™ CardiO2^®^ Gas Exchange Analysis System, MGC Diagnostics^®^, MN, USA), continuously recorded expired air to determine VO_2_. VO_2max_ was determined as the highest 30-s mean VO_2_ coincident with a respiratory exchange ratio equivalent or greater to 1.15.

The day of the study, MAP was measured with a finger photoplethysmograph with a height correcting unit for correction of values (Finometer Pro, Finapres Medical Systems, Amsterdam, The Netherlands or Nexfin, Edwards Lifesciences, Irvine, CA, USA). MCAv was measured with a transcranial Doppler ultrasound (Spencer Technologies, Seattle, WA or Doppler Box, Compumedics DWL USA, San Juan Capistrano, CA) on left temporal window. The MCA was identified using a standardized protocol (38). Once the MCA was identified, a probe was fixed in place with a headband and adhesive ultrasonic gel.

Partial pressure of end-tidal carbon dioxide (P_ET_CO_2_) was recorded beath-by-breath in 48 participants during all the repeated squat-stand through a gas analyzer with fast responding oxygen and carbon dioxide sensors and a pneumotach (Breezesuite, MedGraphics Corp., MN, USA; Ultima™ CardiO2^®^ Gas Exchange Analysis System, MGC Diagnostics^®^, MN, USA; ML206, AD Instruments, Colorado Springs, CO) calibrated to known gas concentrations following manufacturer instructions before each evaluation. The analyzer was adjusted for room temperature, atmospheric pressure and percentage of humidity the day of testing.

All data were simultaneously sampled at 1000 Hz with an analog-to-digital converter (Powerlab 16/30 ML880, AD Instruments) and saved for later analysis using commercially available software (LabChart version 7.1, AD Instruments). There was no delay in recording variables from either transcranial Doppler ultrasound units. In addition, we did not apply any time shift in LabChart. However, to account for established delays in our blood pressure monitoring devices, we applied a −250- and −1,000-ms time shift to ensure Nexfin and Finometer, respectively, and transcranial Doppler ultrasound data were time aligned. Any delays within the data collection units were negated when the structured time-aligned analysis was performed. P_ET_CO_2_ was time-aligned with the other signals.

### Data analysis

To characterize the cerebral pressure-flow relationship in response to transient increases and decreases in MAP, we calculated a time-adjusted ratio between MCAv and MAP changes for each squat-stand transition in each MAP direction. To do so, absolute changes in MCAv (ΔMCAv_T_) and MAP (ΔMAP_T_) for each increase (from minimum to maximum) and decrease (from maximum to minimum) were calculated and adjusted for time intervals when the MCAv and MAP transition took place. Specifically, we divided the changes by the duration of the transition (i.e. time interval between minimum to maximum or between maximum to minimum). Then, we averaged ΔMCAv_T_/ΔMAP_T_ for each individual’s repeated squat-stands over the 5-min recording period and for each frequency.

ΔMCAv_T_/ΔMAP_T_ during transient decreases in MAP was calculated between a maximum and the following minimum as follows (Figure 1):

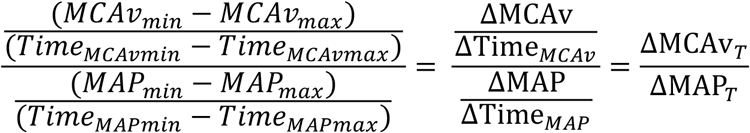

**Figure 1.**
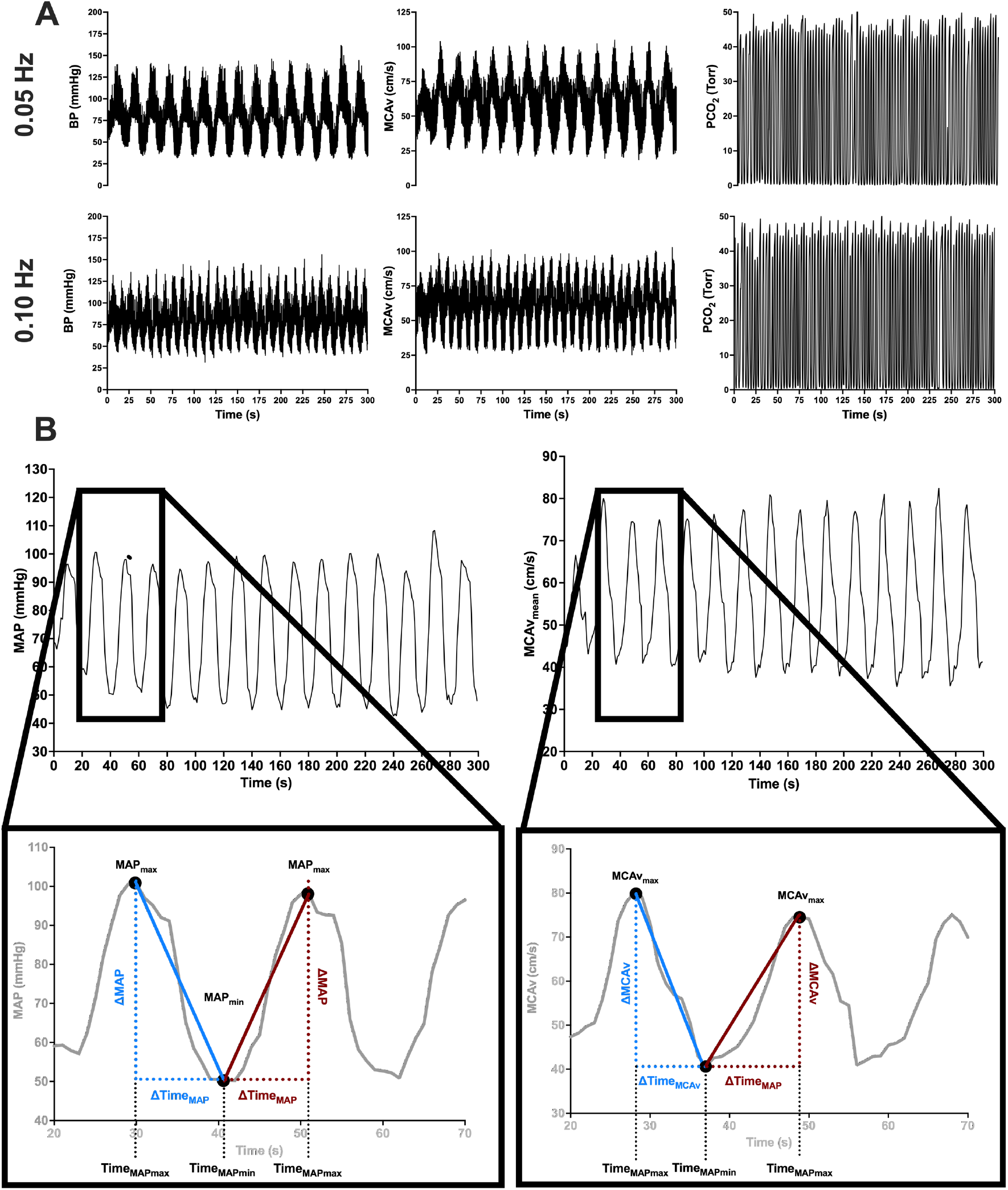
Panel A represents raw tracings of blood pressure (BP), middle cerebral artery blood velocity (MCAv) and partial pressure of carbon dioxide (PCO_2_) at 0.05 and 0.10 Hz repeated squat-stand. Panel B depicts schematic tracings of mean arterial pressure (MAP) and middle cerebral artery mean blood velocity (MCAv) during a repeated squat-stands performed at 0.05 Hz. Enlarged regions depict how the variables were used to calculate ΔMCAv_T_/ΔMAP_T_. Blue color indicates variables used to calculate ΔMCAv_T_/ΔMAP_T_ during acute decreases in MAP. Red color indicates variables used to calculate ΔMCAv_T_/ΔMAP_T_ during acute increases in MAP.

ΔMCAv_T_/ΔMAP_T_ during transient increases in MAP was calculated between a minimum and the following maximum as follows (Figure 1):

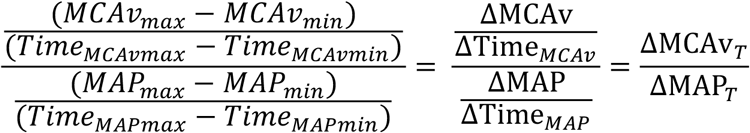

### Statistical analysis

The normality of distributions was tested using Shapiro-Wilk tests. If distributions were normal, a paired t-test was used. If not, the Wilcoxon matched pairs signed rank test was used to compare variables between transient increases and decreases for each frequency. To test the influence of time on ΔMCAv_T_/ΔMAP_T_ over the 5-min recording, a nonparametric repeated-measures Friedman test was performed. Since this test does not support missing values, all participants with missing values for one or more transitions were excluded. Considering the impact of cardiorespiratory fitness and age on the cerebral pressure-flow relationship (4, 5, 22, 24), correlations between ΔMCAv_T_/ΔMAP_T_ and age or VO_2_max were determined using Spearman’s rho correlations. P values < 0.05 were considered statistically significant.

## Results

A total of 74 participants were included in this analysis (9 women, 65 men). For the 0.05 Hz analysis, 6 participants were excluded due to inconsistency of MAP and MCAv oscillations (n=5) and poor recordings of the Nexfin (n=1). For the 0.10 Hz analysis, 3 participants were excluded because squats were not completed (n=1) and poor recordings of the Nexfin (n=2). Final samples for 0.05 Hz and 0.10 Hz repeated squat-stands were 68 and 71, respectively. VO_2_max was determined in 53 participants. Baseline characteristics as well as resting hemodynamic and cerebrovascular seated baseline values are shown in Table 1. Seated baseline MAP and MCAv were missing for 4 participants.

**Table 1.** Baseline characteristics and resting seated values.

| Baseline characteristics |  |
| --- | --- |
| n | 74 |
| Age (years) | 32 ± 13 (20 – 74) |
| Height (m) | 1.75 ± 0.09 (1.55 – 1.94) |
| Body weight (kg) | 75.5 ± 12.8 (53.0 – 115.0) |
| Body mass index (kg/m<sup>2</sup>) | 24.5 ± 3.2 (19.0 – 36.7) |
| Maximal oxygen consumption (ml·kg<sup>-1</sup>·min<sup>-1</sup>) | 46.9 ± 9.6 (23.5 – 65.4) * |
| Resting values |  |
| Mean arterial pressure (mmHg) | 90 ± 13 (60 – 122) <sup>§</sup> |
| Middle cerebral artery mean blood velocity (cm·s<sup>-1</sup>) | 62 ± 13 (34 – 100) <sup>§</sup> |
| End-tidal carbon dioxide partial pressure (Torr) | 38 ± 4 (29 – 47) <sup>†</sup> |

Data are mean ± SD (ranges).
\* n = 53, <sup>§</sup> n = 70, <sup>†</sup> n = 48

Absolute changes in MAP during acute increases in MAP were +43 ± 15 mmHg (0.05 Hz) and +37 ± 14 mmHg (0.10 Hz), whereas they were −43 ± 15 mmHg (0.05 Hz) and −37 ± 14 mmHg (0.10 Hz) during acute decreases in MAP. Absolute changes in MCAv during acute increases in MAP were +36 ± 14 cm·s^-1^ (0.05 Hz) and +32 ± 11 cm·s^-1^ (0.10 Hz), whereas they were −35 ± 14 cm·s^-1^ (0.05 Hz) and −32 ± 11 cm·s^-1^ (0.10 Hz) during acute decreases in MAP. Averaged P_ET_CO_2_ during the 5-min squat-stand repetitions were not different between frequency (0.05 Hz: 38.7 ± 3.8 Torr, 0.10 Hz: 38.2 ± 3.4 Torr; p = 0.0916) and similar to baseline resting values (Table 1).

At 0.05 Hz, averaged time intervals for MAP and MCAv changes were shorter during acute MAP increases than MAP decreases (MAP: 9.0 ± 1.0 vs. 11.0 ± 1.0 sec; p < 0.0001; MCAv: 9.3 ± 1.5 vs. 10.7 ± 1.4 cm·s^-1^; p < 0.0001). At 0.10 Hz, averaged time intervals for MAP and MCAv changes were shorter during acute increases than decreases for MAP (4.9 ± 0.4 vs. 5.2 ± 0.4 sec; p = 0.0024), but not for MCAv (5.1 ± 0.5 vs. 4.9 ± 0.5 sec; p = 0.1028).

For 0.05 Hz repeated squat-stands, ΔMCAv_T_/ΔMAP_T_ was similar during acute increases and acute decreases in MAP (0.88 ± 0.28 vs. 0.90 ± 0.23 cm·s^-1^/mmHg; p = 0.2693; Figure 2). For 0.10 Hz repeated squat-stands, ΔMCAv_T_/ΔMAP_T_ was attenuated during acute increases when compared to acute decreases in MAP (0.91 ± 0.34 vs. 1.01 ± 0.44 cm·s^-1^/mmHg; p = 0.0013; Figure 2).

**Figure 2.**
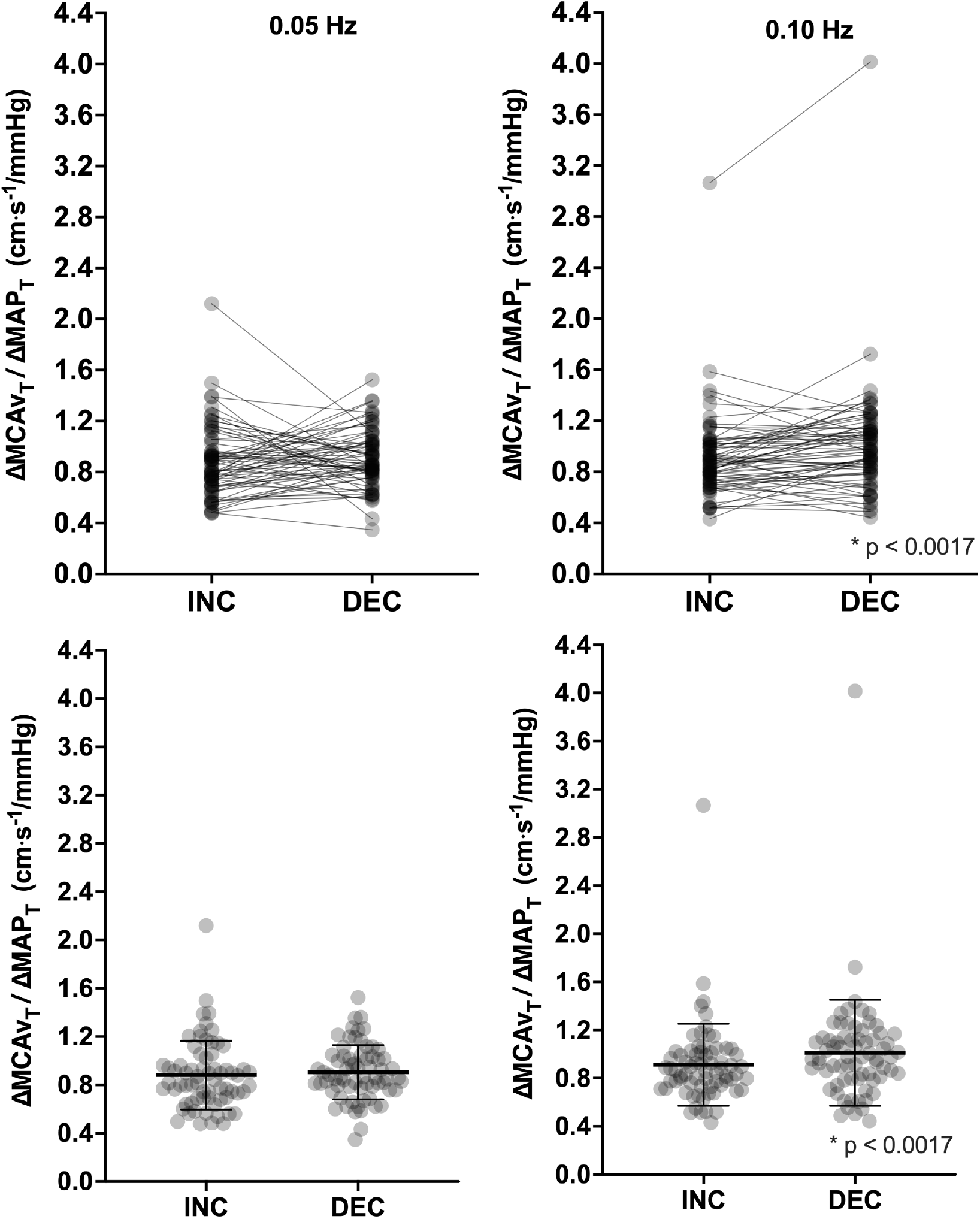
Comparison of the ΔMCAv_T_/ΔMAP_T_ during transient increases (INC) and decreases (DEC) in mean arterial pressure during 0.05 and 0.10 Hz repeated squat-stands. n = 68 for 0.05 Hz, n = 71 for 0.10 Hz. Differences assessed via Wilcoxon matched pairs signed rank test.

For both frequency and MAP direction, time during repeated squat-stands had no effect on ΔMCAv_T_/ΔMAP_T_ (all p> 0.05; Figure 3).

**Figure 3.**
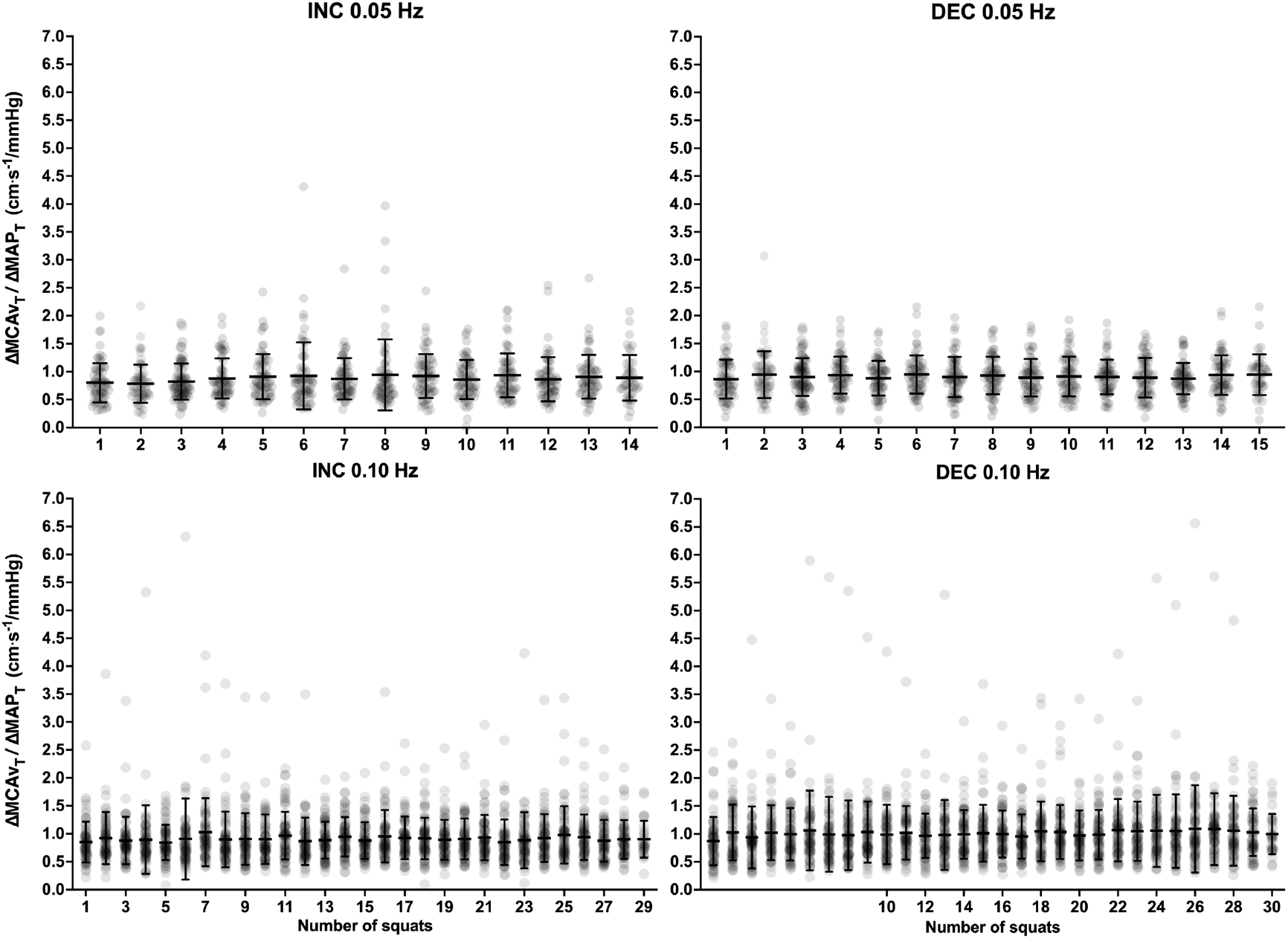
Individual ΔMCAv_T_/ΔMAP_T_ for each squat repetition during acute increases (INC) and decreased (DEC) for 0.05 and 0.10 Hz repeated squat-stands. Numbers on the x-axis represent squat-stand repetitions. For each squat-stand repetition, n = 64 during INC and n = 56 during DEC for 0.05 Hz, n = 58 during INC and n = 59 during DEC for 0.10 Hz. The effect of time was tested using a nonparametric repeated-measures Friedman test.

Cardiorespiratory fitness was correlated to ΔMCAv_T_/ΔMAP_T_ for 0.05 Hz repeated squat-stands during acute MAP increases (ρ = 0.33, p = 0.0244) but not for acute MAP decreases (ρ = 0.21, p = 0.1566). Cardiorespiratory fitness was correlated to ΔMCAv_T_/ΔMAP_T_ for 0.10 Hz repeated squat-stands during both acute increases (ρ = 0.49, p = 0.0003) and decreases (ρ = 0.48, p = 0.0005) in MAP (Figure 4). At 0.05 Hz, age was not correlated to ΔMCAv_T_/ΔMAP_T_ during acute MAP increases (ρ = −0.19, p = 0.1278) nor acute decreases in MAP (ρ = −0.17, p = 0.1581). At 0.10 Hz, age was not correlated to ΔMCAv_T_/ΔMAP_T_ during acute MAP increases (ρ = −0.21, p = 0.0803), but was inversely correlated during acute MAP decreases (ρ = −0.25, p = 0.0346) (Figure 5).

**Figure 4.**
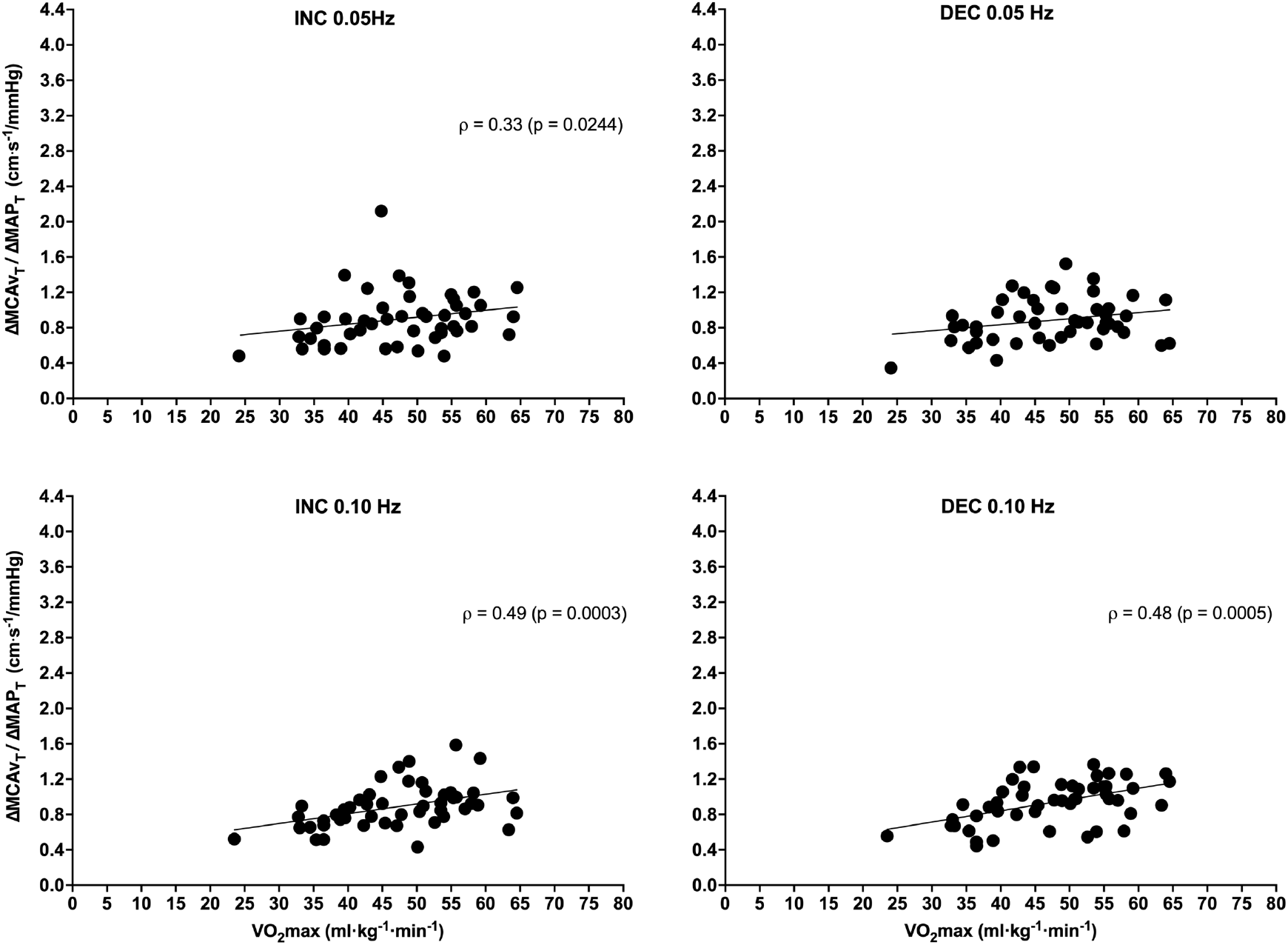
Relationship between VO_2_max and ΔMCAv_T_/ΔMAP_T_ for acute increases (INC) and decreases (DEC) in MAP during repeated squat-stands at 0.05 Hz and 0.10 Hz. n = 47 for 0.05 Hz, n = 50 for 0.10 Hz. Relationships between ΔMCAv_T_/ΔMAP_T_ and cardiorespiratory fitness (VO_2_max) were determined using Spearman’s rho correlations.

**Figure 5.**
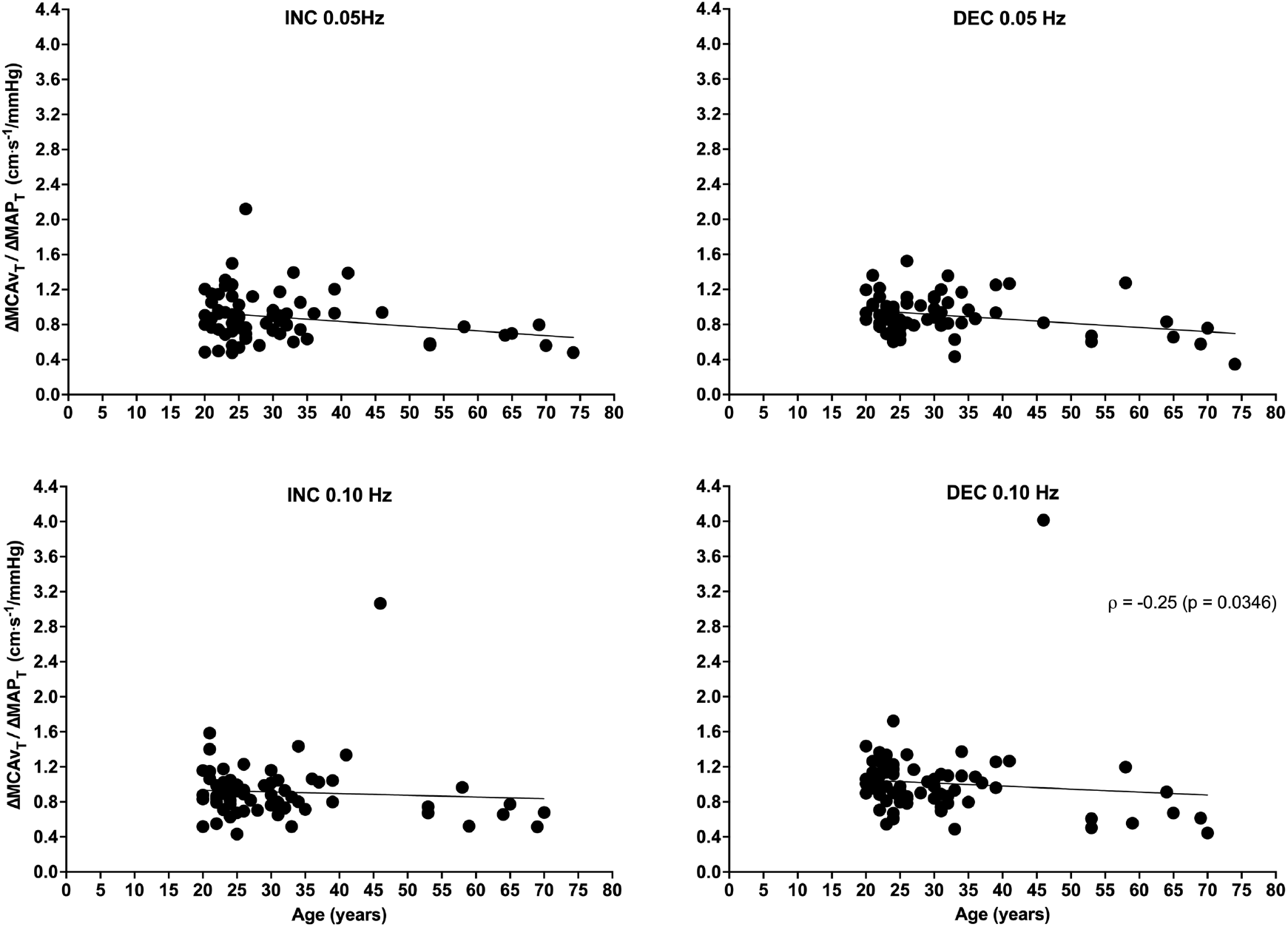
Relationship between age and ΔMCAv_T_/ΔMAP_T_ for acute increases (INC) and decreases (DEC) in MAP during repeated squat-stands at 0.05 Hz and 0.10 Hz. n = 68 for 0.05 Hz, n = 71 for 0.10 Hz. Relationships between ΔMCAv_T_/ΔMAP_T_ and age were determined using Spearman’s rho correlations.

## Discussion

This study demonstrates when MAP is acutely increased from minimum to maximum during repeated squat-stands at 0.10 Hz, the change in MCAv is attenuated compared to when MAP acutely decreases, which is in line with our previous results (8). The novel finding of this study resides in the notion that changes were adjusted for time intervals and calculated on actually induced MAP oscillations, between maximums and minimums *during* the maneuvers, instead of being calculated from a seated baseline value obtained *prior-to* the repeated squat-stands. These results reinforce the evidence of hysteresis in the relationship between MCAv and MAP, indicating cerebral vessels are better adapted to buffer acute increases in MAP at 0.10 Hz, highlighting a probable influence of repeated squat-stand frequency. In addition, our findings convincingly demonstrate the repeated squat-stand model can provide a reliable estimate of the physiological response by averaging several transient responses, considering the physiological stress imposed by each squat-stand repetition leads to comparable ΔMCAv_T_/ΔMAP_T_ (i.e. when the metric for each acute increase in MAP is compared) over the 5-min period, at least in the studied population.

In light of the questions regarding our previous analysis (6), it was important to improve our initial analysis in order to address different concerns, i.e. to avoid the utilization of a separate resting period and take time intervals of MCAv and MAP changes into consideration. To the best of our knowledge, this is the first study to report the presence of a directional sensitivity of the human MCAv-MAP relationship when changes are adjusted for time intervals as well as calculated between each maximums and minimums reached during 0.10 Hz, but not 0.05 Hz, for repeated squat-stands. The results of the present analysis demonstrate the use of actual MAP changes, instead of calculating MAP changes from a separate baseline period, support the presence of a frequency-dependent directional sensitivity of the cerebral pressure-flow relationship in healthy participants. Our study also reinforces the use of the 5-min squat-stand protocol to analyze the MCAv-MAP relationship in the time domain. In fact, as demonstrated in Figure 3, the repetition of the same stimulus over 5 minutes produces valid and stable data over time, reducing the potential influence of outliers.

Although the current study did not aim at examining the mechanisms underlying the asymmetrical sensitivity of the cerebral pressure-flow relationship, multiple hypotheses are discussed in the literature. First, it was shown in an animal model that cerebral sympathetic nervous activity (SNA) recorded in the superior cervical ganglia increases in response to mechanically or drug-induced increases in arterial pressure, but not during transient reduction in arterial pressure (13). These results would indicate cerebral SNA increases in response to an acute elevation in MAP to protect the brain from overperfusion. Interestingly, it is proposed that 0.10 Hz repeated squat-stand frequency might reveal the influence of SNA on human cerebrovascular regulation (20, 40). Therefore, the lower ΔMCAv_T_/ΔMAP_T_ during acute increases in MAP at 0.10 Hz repeated squat-stands could be explained by SNA activation when MAP acutely increases, in order to protect cerebral vessels. However, the impact of cerebral SNA on CBF in humans is still controversial for many reasons (reviewed in (3, 9)). Below frequencies of 0.07 Hz, cerebrovascular regulation is thought to have a myogenic influence (20, 40). Another potential mechanism could therefore be a differing intrinsic myogenic activity when MAP increases than when MAP decreases. Using oscillatory lower body negative pressure and a non-linear approach to quantify dCA, it was demonstrated the utilization of calcium-channel blockade impairs dCA in humans, indicating a role for myogenic mechanisms at frequencies between 0.03 and 0.08 Hz (33). However, these results showed only a global dCA response since the directionality of MAP changes were not considered (33). As MAP direction did not influence ΔMCAv_T_/ΔMAP_T_ at 0.05 Hz repeated squat-stands in the present study, it could be argued the myogenic influence on CBF is similar whether MAP acutely increases or decreases, or that our linear approach masks differences. Recently, Barnes et al. were interested in different components of the cerebrovascular response to repeated acute increases and decreases in MAP during the repeated squat-stand protocol. They demonstrated MAP would be the strongest determinant of CBV when squatting down, overwhelming the myogenic and metabolic components of dCA (6). Overall, these results also make the myogenic mechanism a controversial mechanism to explain the presence of a hysteresis-like pattern in the cerebral pressure-flow relationship (33). The absence of directional sensitivity at 0.05 Hz repeated squat-stands could also be explained by the baroreflex latency (19, 36). In fact, during these slower oscillations, baroreceptors have time to sense pressure loading or unloading and induce appropriate cardiovascular adjustments. This could mitigate the impact of MAP on MCAv over a 10-sec time interval. Finally, a transient increase in MAP could elevate superior vena cava’s resistance, inducing an increase in intracranial pressure and therefore, a reduction of the passive pressure gradient for venous drainage (16, 17), which could represent evidence of the Monro-Kellie doctrine, i.e. a mechanism limiting the amount of CBF increase that is able to occur within the encapsulated skull (39).

The consistency of the ΔMCAv_T_/ΔMAP_T_ over the 5-min repeated squat-stand at both frequencies is indicative the physiological stress induced is stable for both direction and magnitude of acute changes in MAP. The absence of a drift in the values demonstrates the participants performing repeated squat-stands do not get fatigued from the maneuvers and the addition of squat-stand transitions has no influence on the metric, at least in healthy participants.

The relationships between ΔMCAv_T_/ΔMAP_T_ and cardiorespiratory fitness during acute increases and decreases in MAP at 0.10 Hz, and only during acute increases at 0.05 Hz repeated squat-stand protocol, are also interesting observations. These findings suggest a diminished ability to cope with higher frequency MAP oscillations with increasing cardiorespiratory fitness. We have previously demonstrated an association between cardiorespiratory fitness and TFA gain, normalized gain and phase during 0.10 Hz repeated squat-stands in young and healthy fit men (22). TFA gain metrics represent changes in MCAv for a given change in MAP, which makes logical that ΔMCAv_T_/ΔMAP_T_ is also correlated to cardiorespiratory fitness in the current analysis. However, the fact that the current analysis using the ΔMCAv_T_/ΔMAP_T_ metric takes the directionality of MAP changes into account is notable. The reason for an augmented ΔMCAv_T_/ΔMAP_T_ during acute increases and decreases in MAP, (which could be interpreted as a diminished ability to buffer MAP changes in both directions, i.e. a lowered dCA), with increasing cardiorespiratory fitness and whether this is deleterious or beneficial for the brain, is still unknown. Further studies are needed to evaluate the impact of improving cardiorespiratory fitness on the maintenance of the hysteresis-like pattern in the MCAv-MAP relationship.

In the current analysis, age inversely correlates with ΔMCAv_T_/ΔMAP_T_ when MAP acutely decreases during 0.10 Hz repeated squat-stands. These findings contradict results from our previous work (8). Indeed, when calculating relative changes from a seated baseline obtained prior to the repeated squat-stand protocol, age was positively correlated with %ΔMCAv/%ΔMAP when MAP acutely increased for 0.05-Hz repeated squat-stand only. As in our previous study, the correlational analysis included here is exploratory in nature. Statistical power may have been insufficient to address that question. In addition, while we have included a group of participants with a large range in age, only a small number of subjects were above 40 years old, which considerably reduce the strength of this analysis. Further well-designed research aimed at specifically examining the influence of age on the directional sensitivity of the cerebral pressure-flow relationship is thus necessary to address that specific question.

### Perspectives

Overall, our updated analysis for repeated squat-stands is adequate to study the directional sensitivity of the cerebral pressure-flow relationship and optimize the assessment of dCA. It supports the need of a global change in dCA assessment in order to take the direction of MAP into account. It also reinforces the importance of using the repeated squat-stands, at different frequencies, to provide a broader characterization of the cerebral pressure-flow relationship. In addition, since we showed repeated squat-stands induce a comparable change in ΔMCAv_T_/ΔMAP_T_ at each squat-stand transition, it eliminates the questioning as to whether the addition of squat-stands (exercise) would have an impact *per se* on this metric. A next important step of major interest would be to evaluate the reproducibility of this metric over time. It would help determine whether the ΔMCAv_T_/ΔMAP_T_ is reliable over various time frames and if it remains stable over time. Still, further research is needed to examine the possibility of using less repetitions to calculate the ΔMCAv_T_/ΔMAP_T_ metric. Indeed, a diminished number of squat-stand repetitions would be an important methodological advantage to examine for the presence of hysteresis in clinical populations for whom a 5-min period of repeated squat-stands could potentially be difficult to complete.

The directional sensitivity in the cerebral pressure-flow relationship is highly relevant for various physiological (rapid eye movement sleep, exercise), and pathological clinical situations associated with MAP surges. Nonetheless, further research is needed to describe this phenomenon in other populations and clinical conditions. We still do not know whether this hysteresis-like pattern remains present in different physiological and pathological states (arterial/carotid stiffness, uncontrolled systemic hypertension, autonomic dysreflexia, etc.). For example, findings from a recent study using a mouse model of arterial stiffness suggest mice with calcified carotid artery have a disrupted cerebral autoregulation in response to steady-state MAP increases compared to healthy animals (26). The next logical step would therefore be to examine whether there is evidence for hysteresis in the cerebral pressure-flow relationship in human clinical populations and the present study serves as a fundamental basis to further explore the phenomenon. Importantly for the clinician, this directional sensitivity of brain vessels to changes in MAP should also be integrated in the framework to guide its management in the operating room. Finally, considering the link found with cardiorespiratory fitness in the present study, it would be interesting to evaluate the impact of exercise training on the maintenance of the hysteresis-like pattern.

### Methodological considerations

There are some limitations to this study that need to be further discussed. Only healthy men and women were included in this analysis and the results cannot be generalized to clinical populations. Moreover, MCAv was evaluated through transcranial Doppler ultrasound, which is known to be representative of flow only if the diameter of the artery is assumed to remain stable. Changes in P_ET_CO_2_ have been associated with MCA’s diameter variations (23, 37). Although P_ET_CO_2_ was not available for a part of the male participants, there was no difference between P_ET_CO_2_ during baseline and repeated squat-stand for the participants in whom it was measured. In addition, it was previously observed that there was no change in P_ET_CO_2_ between squat and stand phases (27). Therefore, P_ET_CO_2_ had most likely a minor effect on our results. Only a small part of the group was composed of women either taking oral contraceptives, wearing an intrauterine device or being in the days 1-10 of their menstrual cycle. We are not able to determine whether it had an influence on the results and further research is needed to address that issue. Since days 0-10 represents a large time period, it should be shortened in future studies in order to better control for menstrual cycle. Of note, all participants (n=58) included in our previous work (8) were also included in the current analysis. From an analytical standpoint, the two approaches are comparable, as both methods are calculating changes in MCAv and MAP in response to repeated squat-stands. One difference resides in the fact the previous method calculated relative changes from a seated baseline, whereas the revised method calculated actual absolute changes adjusted for time intervals. This being acknowledged, the main difference resides in the *baseline* value used to calculate these changes in MCAv and MAP. In the original approach, we have calculated a *baseline* including averaged MCAv and MAP during a steady-state rest. However, this *baseline* with participants in a seated position was measured during a distinct recording period prior to the start of the series of repeated squat-stands. Accordingly, from a physiological standpoint, it makes definitely more sense to calculate changes in MCAv and MAP on actually induced MAP oscillations, between maximums and minimums during the repeated squat-stands, rather than being calculated from a seated baseline value obtained during a distinct period prior to the repeated squat-stands. We consider the updated analytical method proposed in the current work to be the most valid option to examine the directional sensitivity of the cerebral pressure-flow relationship with the repeated squat-stand model since our calculated metric cannot be influenced by a separate resting steady-state baseline. However, the two analytical methods were not directly compared, as our main goal of the current investigation was to examine whether the hysteresis-like pattern in the cerebral pressure-flow relationship was present with the updated analytical method. A direct comparison would not provide additional meaningful information.

## Conclusion

Our findings indicate that this novel analytical method supports the use of the repeated squat-stand model to examine the directional sensitivity of the cerebral pressure-flow relationship, during actual changes in MAP and cerebral blood velocity, without the need for a separate resting baseline. Moreover, it shows this maneuver induces stable and robust physiological cyclic changes and allows for interindividual variability analysis. Importantly, the results contribute to the importance of considering the direction of MAP changes when evaluating dCA and builds upon the prior rate-of-regulation and autoregulatory index measures.

## Acknowledgments

We would like to thank Myriam Paquette, Olivier Le Blanc, Kevan Rahimaly, Sarah Imhoff, Simon Malenfant and Audrey Drapeau for their assistance in data collection.

